# High-resolution retrospective single cell lineage tracing with mutable homopolymers

**DOI:** 10.64898/2026.03.10.709901

**Authors:** Pin-Chung Cheng, Dmitrii Kamenev, Polina Kameneva, Conor Fitzpatrick, Igor Adameyko, Peter V Kharchenko, Kun Zhang

## Abstract

Retrospective reconstruction of cell lineages from somatic mutations holds substantial promise for understanding development, tissue homeostasis, and disease. Yet, current approaches are often constrained by low marker density, technical noise, and incomplete molecular readouts that limit the resolution and accuracy of inferred lineages. Here, we present RETrace2, a single-cell dual-omic method for simultaneous lineage tracing and cell-type identification. By identifying and targeting highly mutable homopolymers, which we show are approximately twice as informative as conventional microsatellite repeats, combined with technical improvements to increase marker coverage and reduce experimental errors of reading and amplifying homopolymers, RETrace2 can trace lineage continuously at an estimated resolution of fewer than five cell divisions in microsatellite-unstable organisms. Simultaneously, sparse methylation profiles were obtained from the same cells for cell-type identification. We validated the method using *in vitro* ground-truth models and demonstrated its utility *in vivo* by reconstructing multi-organ lineage trees from the brain, kidney and liver of *Msh2*-deficient mice.

## Introduction

The ability to trace the lineage of individual cells throughout the lifespan is crucial for understanding how a single fertilized egg develops into a complex organism, and how chronic diseases arise and progress^1,2^. In model organisms, prospective lineage tracing using genetically engineered barcodes has been instrumental in mapping developmental pathways^3–8^. However, traditional prospective methods provide only a static snapshot of clonal history initiated at discrete time points. While newer CRISPR-based approaches enable cumulative recording, they are constrained by target site saturation, preventing the continuous tracking of cell divisions across an entire lifespan. Furthermore, they often require complex genetic engineering and technical designs that are challenging to implement and generally inapplicable to human studies.

Retrospective lineage tracing methods, which infer the history of cell divisions from naturally occurring somatic mutations, offer a powerful alternative^9–11^. Because somatic mutations accumulate with every cell division, they create a continuous, permanent log of a cell’s history. This allows for the reconstruction of high-resolution cellular lineage trees without preconceived genetic labeling or complex experimental setups, making the approach uniquely scalable across different organisms and tissues. To date, various somatic mutations have been explored as lineage barcodes, including mitochondrial DNA mutations^12–16^, DNA methylation epimutations^17,18^, single nucleotide variations (SNVs)^19–27^, and microsatellite instability^28–32^. However, many of these popular markers have significant limitations. For instance, mitochondrial DNA (mtDNA) variants exhibit complex dynamics such as genetic drift and rapid turnover, making it challenging to reconstruct high-resolution cellular phylogenies^33,34^. DNA methylation epimutations can be unstable and confounded by changes in cell type or state^35^, posing challenges for analysis in heterogeneous tissues. While SNVs represent stable genomic markers, their relatively low mutation rate means that accurately profiling them genome-wide remains prohibitively expensive for large-scale single-cell studies.

Among these options, microsatellites, short tandem repeats of DNA, are particularly promising. They represent more stable, permanent lineage markers while possessing 1,000 to 10,000 times higher mutation rates than SNVs. Their high mutation rate stems from polymerase slippage during DNA replication^36^, estimated at approximately 10^−5^ mutations per locus per cell division in stable cells, and increasing to 10^−3^ or higher in microsatellite-unstable systems^37,38^. We previously developed RETrace v1, providing a proof-of-concept for using microsatellites for lineage tracing together with methylation profiling. However, its resolution was limited by the limited somatic mutation rates of di-to hexa-nucleotide repeats, as well as low numbers of informative markers captured per cell^29^.

Here, we report RETrace2, a substantially improved dual-omic method that utilizes highly mutable mononucleotide microsatellites (homopolymers) for lineage construction alongside DNA methylation for cell typing. We demonstrate that homopolymers are approximately 1.9-fold more informative as lineage markers compared to other repeat types. To demonstrate this potential, we systematically optimized the experimental protocol, achieving a ∼21-fold increase in the median number of markers captured per cell and a ∼98-fold increase in shared markers per cell pair compared to RETrace v1.

Furthermore, by systematically addressing the inherent susceptibility of homopolymers to experimental artifacts, we greatly reduced errors in both amplification and sequencing. We validated the resolution and accuracy of RETrace2 *in vitro* and demonstrated *in vivo* cell lineage construction of brain, kidney, and liver tissues from *Msh2*-mutant mice, establishing a framework for continuous, high-resolution, multi-organ lineage tracing.

## Results

### Homopolymers as superior lineage tracing markers

RETrace2 utilizes a dual-omic library preparation strategy optimized to simultaneously capture lineage and cell type information from the same single cells (Fig. 1a). Single cells or nuclei are isolated via FACS into individual wells containing a lysis buffer. The genomic DNA is subjected to a dual-restriction enzyme digestion: MseI enriches for A/T-rich microsatellite regions, while MspI targets C/G-rich regions for methylation profiling. Following adapter ligation, we employed a single primer PCR step. This critical step performs linear amplification of the original template, generating high-fidelity copies of the initial genetic material before exponential amplification. The library is subsequently split: one aliquot is processed for hybridization capture using a custom biotinylated probe panel to selectively enrich for the informative microsatellite targets, while the second undergoes bisulfite conversion for methylation profiling. Finally, both libraries are pooled and sequenced, allowing us to reconstruct the lineage of each cell while simultaneously determining its cellular identity.

**Fig. 1.**
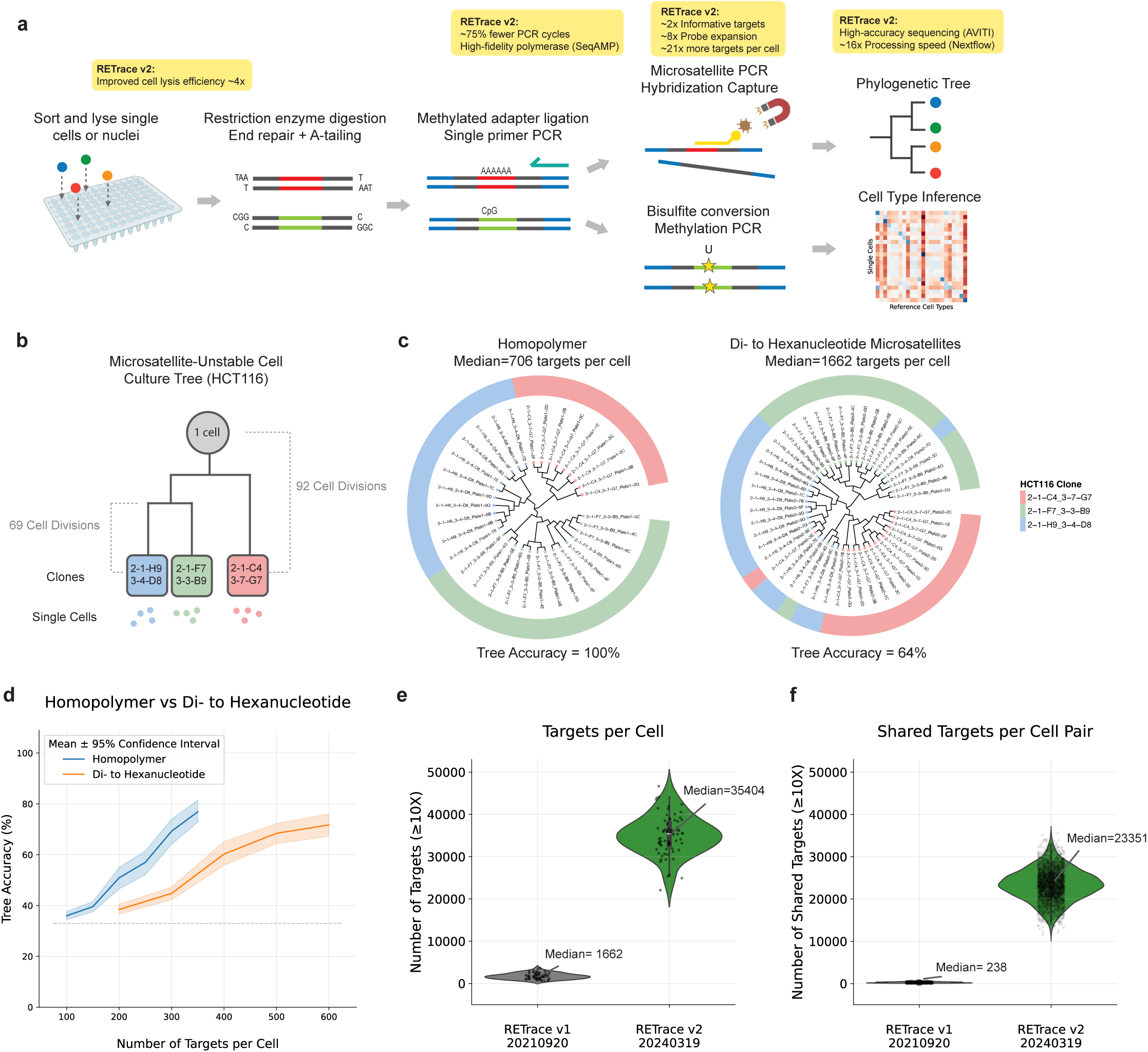
RETrace2 efficiently captures highly informative homopolymer lineage markers. **a,** Schematic of the RETrace2 workflow. Created with BioRender. **b,** HCT116 ground-truth cell culture model for benchmarking microsatellite-unstable cells. Dashed lines indicate number of cell divisions from root/node to clone. **c,** Reconstructed HCT116 phylogenetic tree with homopolymer (n=34 cells) or di- to hexanucleotide microsatellite targets (n=49 cells). **d,** Tree accuracy as a function of the number of targets. Lines show mean accuracy from 30 random downsampling iterations; shade represents the 95% confidence interval. The dashed line indicates random accuracy (33%). **e,** Number of targets captured per cell for RETrace v1 (20210920 di to hexanucleotide library; n=49) and v2 (20240319 library; n=72). Each dot represents a single cell. **f,** Number of shared targets per cell pair for RETrace v1 (n=1,176 pairs) and v2 (n=2,556 pairs).

A key limitation of our previous method, RETrace v1, was the reliance on di- to hexa-nucleotide microsatellites, which have relatively lower mutation rates. To improve lineage tracing resolution, we hypothesized that mononucleotide microsatellites, or homopolymers, would serve as more informative markers due to their higher mutability^39^. We benchmarked this hypothesis by applying two distinct probe sets, one targeting homopolymers and another targeting di- to hexa-nucleotide repeats, to the same microsatellite-unstable cell culture model (HCT116) with a known ground-truth lineage (Fig. 1b). Phylogenetic trees reconstructed from the homopolymer markers achieved 100% accuracy, correctly clustering all cells according to their clonal origin. In contrast, the tree reconstructed from di- to hexa-nucleotide markers achieved only 64% accuracy, despite capturing a higher median number of targets per cell in this comparison (706 homopolymers vs. 1,662 di- to hexa-nucleotides) (Fig. 1c). To estimate the information content of these markers, we performed a computational down-sampling analysis. We found that significantly fewer homopolymer markers were sufficient to achieve the same level of tree accuracy as the conventional repeats (Fig. 1d). Linear regression analysis revealed that the accuracy slope for homopolymers was approximately 1.9-fold steeper than for di- to hexa-nucleotides (0.171 vs. 0.090 accuracy increase per target; ANCOVA interaction *p* < 0.001, (Extended Data Fig. 1a), confirming that homopolymers are more informative markers for lineage reconstruction.

We next sought to maximize the number of targetable loci per cell. We validated that shorter, highly abundant 10–14 bp homopolymers were similarly informative to longer 15–50 bp repeats (Extended Data Fig. 1b, c). This allowed us to expand our probe set approximately 8-fold by including these abundant short repeats (Extended Data Fig. 1d, e). In parallel, we comprehensively optimized the experimental protocol to address the low capture efficiency of RETrace v1. We developed an improved cell lysis buffer that increased initial DNA yield by four-fold (Extended Data Fig. 1f) and implemented a new probe design with increased targeting area and universal blockers to enhance hybridization efficiency (Extended Data Fig. 1g). Collectively, these enhancements resulted in a ∼21-fold increase in the median number of markers captured per cell (1,662 in v1 vs. 35,404 in v2) (Fig. 1e) and, crucially, a ∼98-fold increase in the number of shared markers per cell pair for lineage construction (238 in v1 vs. 23,351 in v2) (Fig. 1f).

### Reducing homopolymer artifacts

A technical challenge in employing homopolymers is their inherent susceptibility to experimental errors. To quantitatively assess and mitigate these errors, we developed a synthetic oligonucleotide model to benchmark fidelity (Fig. 2a). This model comprised a pool of five oligonucleotides with distinct microsatellite sequences: three homopolymers (15xA, 20xA, 30xA) and two dinucleotide repeats (10xAC, 15xAC) (Supplementary Table 2). These oligonucleotides were chemically synthesized, with each molecule tagged with a Unique Molecule Identifier (UMI) for deriving low-error consensus sequences. By amplifying this library with varying PCR cycles and sequencing the products, we calculated a per-UMI accuracy by comparing the microsatellite sequence from each individual read (Read 1) to the consensus sequence derived from all reads sharing the same UMI.

**Fig. 2.**
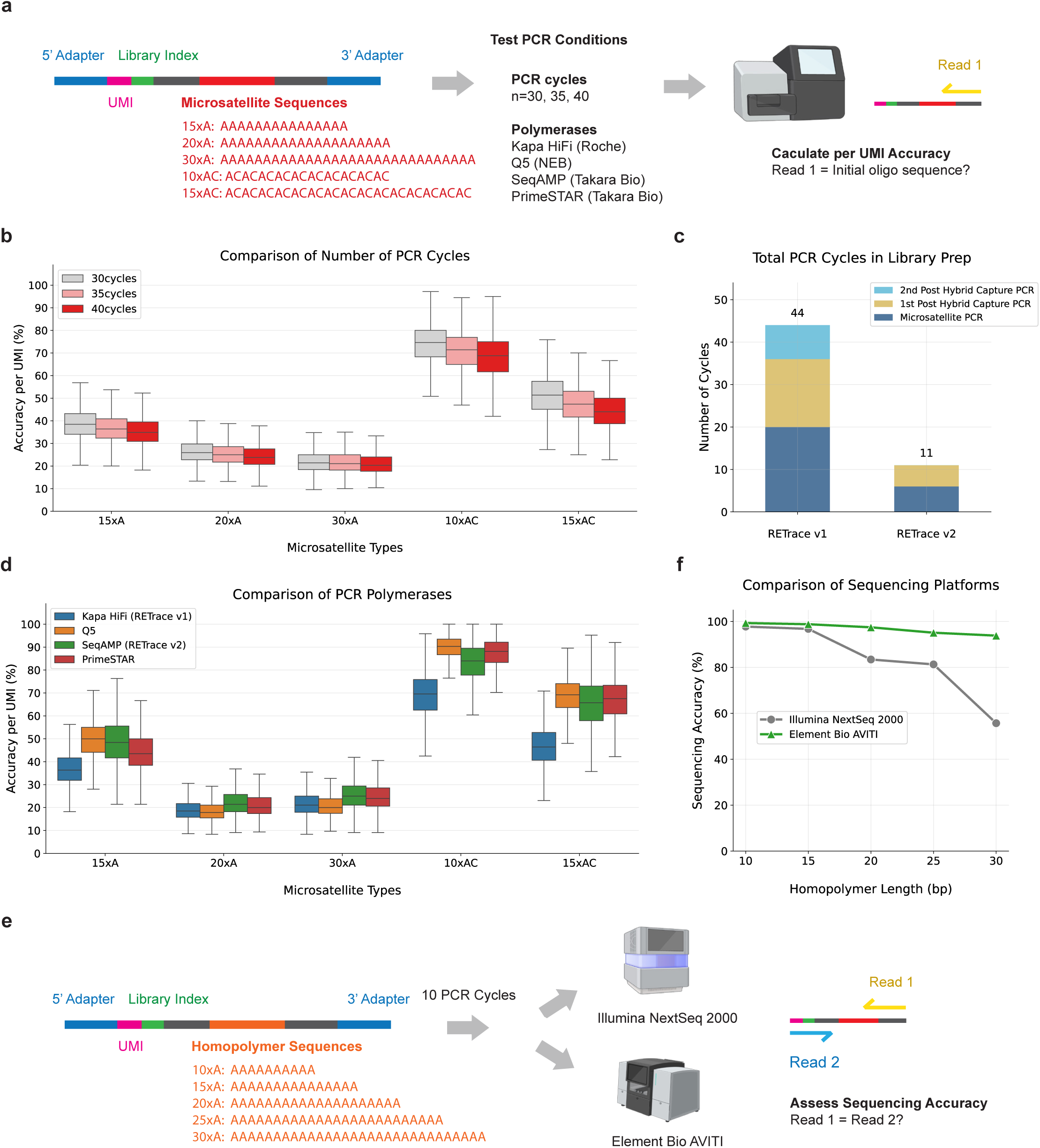
Systematic reduction of homopolymer amplification and sequencing artifacts. **a,** Schematic of the synthetic oligonucleotide model used to measure PCR errors. **b,** Accuracy per unique molecular identifier (UMI) versus the number of PCR cycles. The decrease in accuracy with more cycles was statistically significant for all tested repeat types (one-way ANOVA, all P < 10-38). **c,** Total number of PCR cycles used in RETrace v1 and v2 library preparation. **d,** Accuracy per UMI for multiple DNA polymerases. SeqAMP and PrimeSTAR showed significantly higher accuracy across all microsatellite repeat types compared to Kapa HiFi (the polymerase used in RETrace v1). Q5 showed higher accuracy for 15xA homopolymer and dinucleotide repeats but not for larger homopolymers (Mann-Whitney U test with Bonferroni correction, P < 10-6 for all significant comparisons). **e,** Workflow to measure homopolymer sequencing accuracy. **f,** Comparison of sequencing platform accuracy for homopolymers of different lengths.

We confirmed that accuracy decreased with more PCR amplification cycles (Fig. 2b). Motivated by this finding, and leveraging the improved capture efficiency of the RETrace2 protocol, we optimized the workflow to achieve a ∼75% reduction in total cycles (from 44 in v1 to 11 in v2) (Fig. 2c). In addition, as prior studies have shown, different polymerases exhibit variable stutter errors^40^. We used this oligo model to benchmark multiple high-fidelity polymerases, and identified SeqAMP as a more accurate and robust enzyme for amplification (Fig. 2d).

Finally, in addition to reducing errors prorogated during library preparation, we also sought to minimize errors arising from the sequencing process itself. We used the same oligonucleotide model, focusing on homopolymers between 10–30 bp to match our RETrace v2 probe targets. After a 10-cycle PCR amplification, we split the library for paired-end sequencing on the Illumina NextSeq 2000 and Element Bio AVITI platforms (Fig. 2e). By comparing whether Read 1 and Read 2 sequences matched for each molecule, we could estimate errors generated during sequencing. While the Illumina platform performed well on shorter 10 bp and 15 bp homopolymers, its accuracy dropped significantly as length increased, falling to 55.7% at 30 bp. In contrast, the Element Bio AVITI platform maintained a high accuracy across the full range, retaining 93.8% accuracy at 30 bp (Fig. 2f), a trend consistent with AVITI’s published results^41^. The adoption of the AVITI platform in RETrace2 is therefore beneficial for increasing the accuracy for the longer homopolymer targets.

### Benchmarking on an in vitro cell culture model

To rigorously benchmark the performance of RETrace2, we utilized a cell culture model with a known ground-truth lineage, a strategy established for validating lineage tracing methods^28,29^. This model is constructed by using FACS to deposit single cells into individual wells of a 96-well plate. These single cells are clonally expanded to a population of approximately two million cells over an estimated 21–23 cell divisions (Fig. 3a). A first-generation clone can then be used as the source for subsequent rounds of single-cell sorting and expansion to create a multi-generational tree with a defined branching structure and known relationships between all clones.

**Fig. 3.**
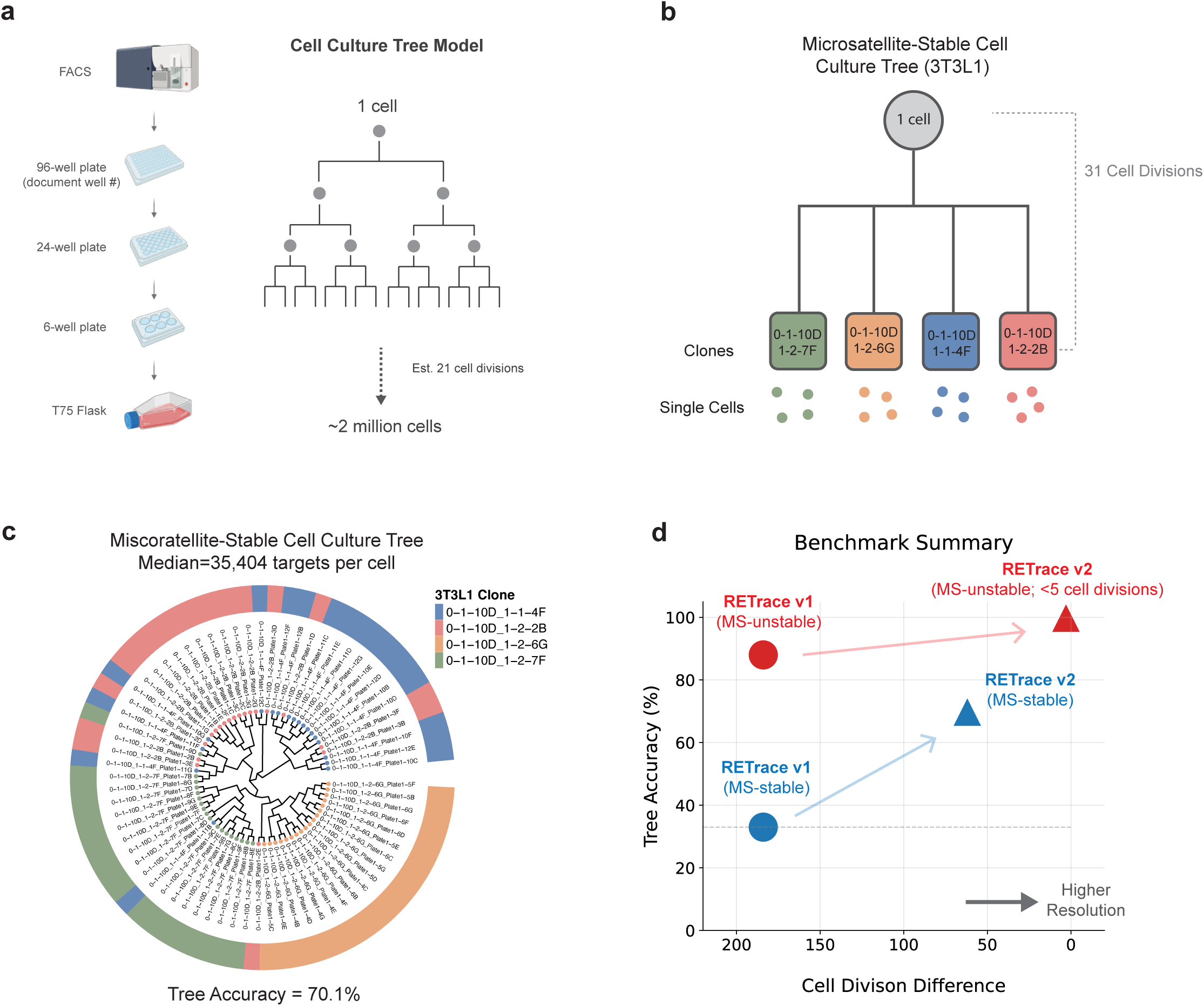
Benchmarking RETrace2 resolution in microsatellite-stable and unstable cell culture models. **a,** Schematic of constructing one generation cell culture model with a known ground truth lineage. Created with BioRender. **b,** 3T3L1 ground-truth cell culture model for benchmarking microsatellite-stable cells. **c,** Reconstructed 3T3L1 phylogenetic tree (n=72) with RETrace2. **d,** Summary of in vitro benchmarking, comparing accuracy and resolution of RETrace v1 (circle) and RETrace2 (triangle) in microsatellite-unstable (red) and microsatellite-stable (blue) cells. Resolution is defined by the number of cell divisions separating clones.

In our previous work, RETrace v1 failed to resolve lineage in microsatellite-stable cells, yielding a baseline random accuracy of ∼33%. To determine if our optimizations could overcome this limitation, we assessed the accuracy of RETrace2 in a mouse microsatellite-stable (3T3L1) cell line, representing a “normal” tissue context with low background mutation rates. We constructed a ground-truth model of four 3T3L1 clones separated by an estimate of 62 cell divisions (Fig. 3b). RETrace2, capturing a median of 35,404 targets per cell, successfully reconstructed the lineage tree with 70.1% accuracy (Fig. 3c). This result establishes that RETrace2 is a viable tool for tracing lineage in normal, non-cancerous tissues where somatic mutations are sparse.

Furthermore, we can extrapolate this performance to cells or tissues with microsatellite instability. Our initial benchmark in HCT116 cells achieved 100% accuracy at 168 divisions with only 706 homopolymers (Fig. 1c). As RETrace2 now captures nearly 50-fold more informative markers per cell (35,404 vs 706) and 98-fold more shared markers per cell pair (23,351 vs 237), we project a resolution of fewer than five cell divisions in microsatellite-unstable systems (Fig. 3d). However, direct validation of this resolution is currently limited by the difficulty of generating ground-truth models with such high temporal precision.

### Clonal analysis in microsatellite instability mouse model

To demonstrate the utility of RETrace2 in a complex *in vivo* setting, we utilized a mouse model with a knockout of Msh2^42,43^, a critical DNA mismatch repair gene. The absence of *Msh2* leads to a hypermutable state known as microsatellite instability (MSI), which accelerates the accumulation of microsatellite mutations and provides an ideal system for high-resolution lineage reconstruction. We generated homozygous *Msh2* knockout (KO/KO) mice and confirmed their microsatellite-unstable status across brain, kidney, and liver tissues (Extended Data Fig. 2a). Importantly, these mice developed normally with no visible tumors in the collected tissues at 6.5 weeks of age, establishing this as an appropriate model for studying developmental lineage rather than just tumorigenesis.

We applied RETrace2 to simultaneously profile 152 single cells from seven distinct tissue regions across the brain (anterior/posterior cortex), kidney (left/right cortex), and liver (left/median/right lobes) of a single 6.5-week-old female mouse (Fig. 4a).

**Fig. 4.**
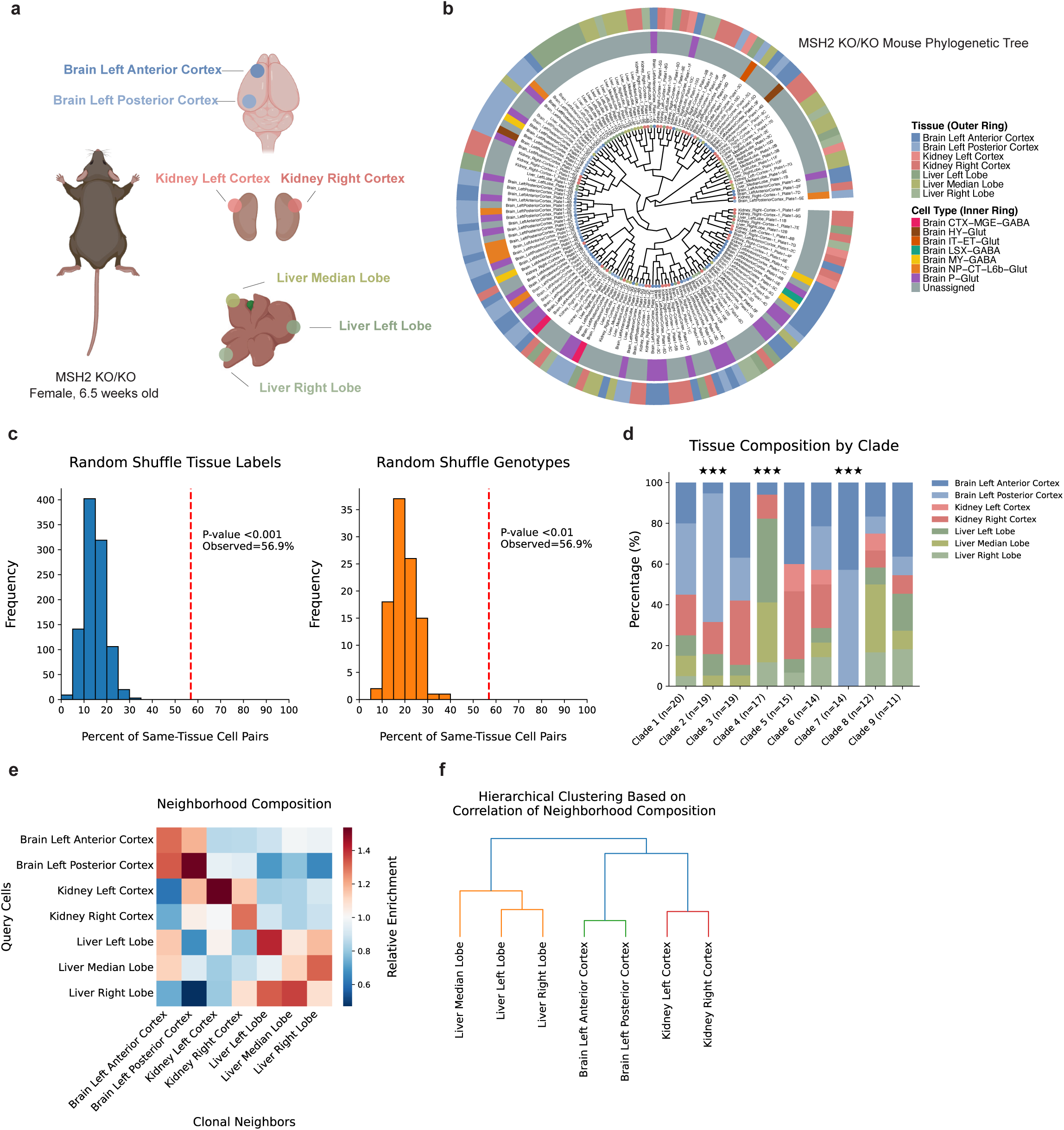
In vivo clonal analysis of multi-organ development in a microsatellite instability mouse model. **a,** Schematic showing the seven tissue regions dissected from a 6.5-week-old female mouse across three organs: brain, kidney, and liver. Created with Biorender. **b,** Reconstructed phylogenetic tree of 152 single cells. The outer ring is colored by tissue of origin, and the inner ring is colored by inferred cell type. **c,** Permutation tests evaluating the statistical significance of tissue-specific clustering. The observed percentage of closely related cell pairs from the same tissue (red dashed line) is significantly higher than null distributions generated by shuffling tissue labels (top, p < 0.001) or microsatellite genotypes (bottom, p < 0.01). **d,** Tissue composition of the nine largest clades (10-20 cells each). Stars indicate significant tissue enrichment (hypergeometric test with FDR correction): *** = strong bias (FDR < 0.05, FC ≥ 2.0); ** = moderate bias (FDR < 0.1, FC ≥ 1.5); * = weak bias (FDR < 0.2, FC > 1.0). **e,** K-nearest neighbor (KNN) analysis (k=15) of clonal relationships. A heatmap showing the relative enrichment of clonal neighbors**. f,** Hierarchical clustering of tissues based on the correlation of their neighborhood composition profiles.

Sequencing yielded high-quality libraries with a median of 11.0 million reads per cell, resulting in a median of 18,983 targets covered at ≥ 10X depth (Extended Data Fig. 2b). The alignment rates for the microsatellite library were consistently high across all tissues, with a mean of 99.87% (Extended Data Fig. 2c).

Using these data, we reconstructed a high-resolution phylogenetic tree (Fig. 4b). The topology is characterized by a marked asymmetry at the root followed by complex, intermixed clonal structures. This overall structure aligns with findings from the prospective LoxCode barcoding study showing that organ development is polyclonal, with some early epiblast clones contributing broadly across tissues while others become biased toward specific lineages^8^. To statistically validate that the tree topology reflects a true biological signal, we performed two permutation tests. We defined the test statistic as the percentage of “sibling tips“—cells sharing the same immediate parent node—that originated from the same tissue (observed value: 56.9%). We compared this to null distributions generated by (1) randomly shuffling tissue labels (p < 0.001) and (2) shuffling the microsatellite genotypes themselves (p < 0.01) (Fig. 4c). In both cases, the observed clustering was significantly higher than random expectation, confirming that the reconstructed tree captures a strong, non-random biological signal.

To assess lineage commitment, we identified nine major clades (subtrees containing 10–20 cells) and evaluated them for tissue-specific enrichment. A hypergeometric test revealed that one-third of these clades showed statistically significant bias toward a single tissue region, while the remaining two-thirds were classified as multipotent, containing cells from multiple tissues without significant enrichment (Fig. 4d). This confirms that tissue-restricted and broad-potential progenitors coexist during the early stages of development.

Finally, to further characterize lineage relationships, we analyzed the clonal neighborhood for each cell using a K-nearest neighbor (KNN) approach (k=15) based on divergence-adjusted genetic distances. A normalized neighborhood composition heatmap showed a pronounced diagonal enrichment pattern, confirming that cells from a given tissue are clonally closer to one another (Fig. 4e). Hierarchical clustering based on the similarity of these neighborhood profiles further grouped tissues by their organ of origin (brain, kidney, and liver), reflecting organ-level clonal relationships (Fig. 4f).

### Inferring cell type from sparse methylation data

The dual-omic nature of RETrace2 allows for the simultaneous capture of cell lineage and cell identity. The methylation profiling workflow is adapted from the single-cell reduced representation bisulfite sequencing (scRRBS) protocol^44^. From the 152 mouse cells that passed quality control, we generated a median of 1.42 million reads per cell with a median alignment rate of 39.97% (Extended Data Fig. 3a, b). Key quality metrics include a high median bisulfite conversion rate of 97.70% and global methylation rates of 36.58% in the CG context and 4.18% in the CH context (Extended Data Fig. 3c, d). We captured a median of 10,770 CpG sites per cell (Fig. 5a), which provides sufficient information to accurately infer cell type given an established high-quality reference.

**Fig. 5.**
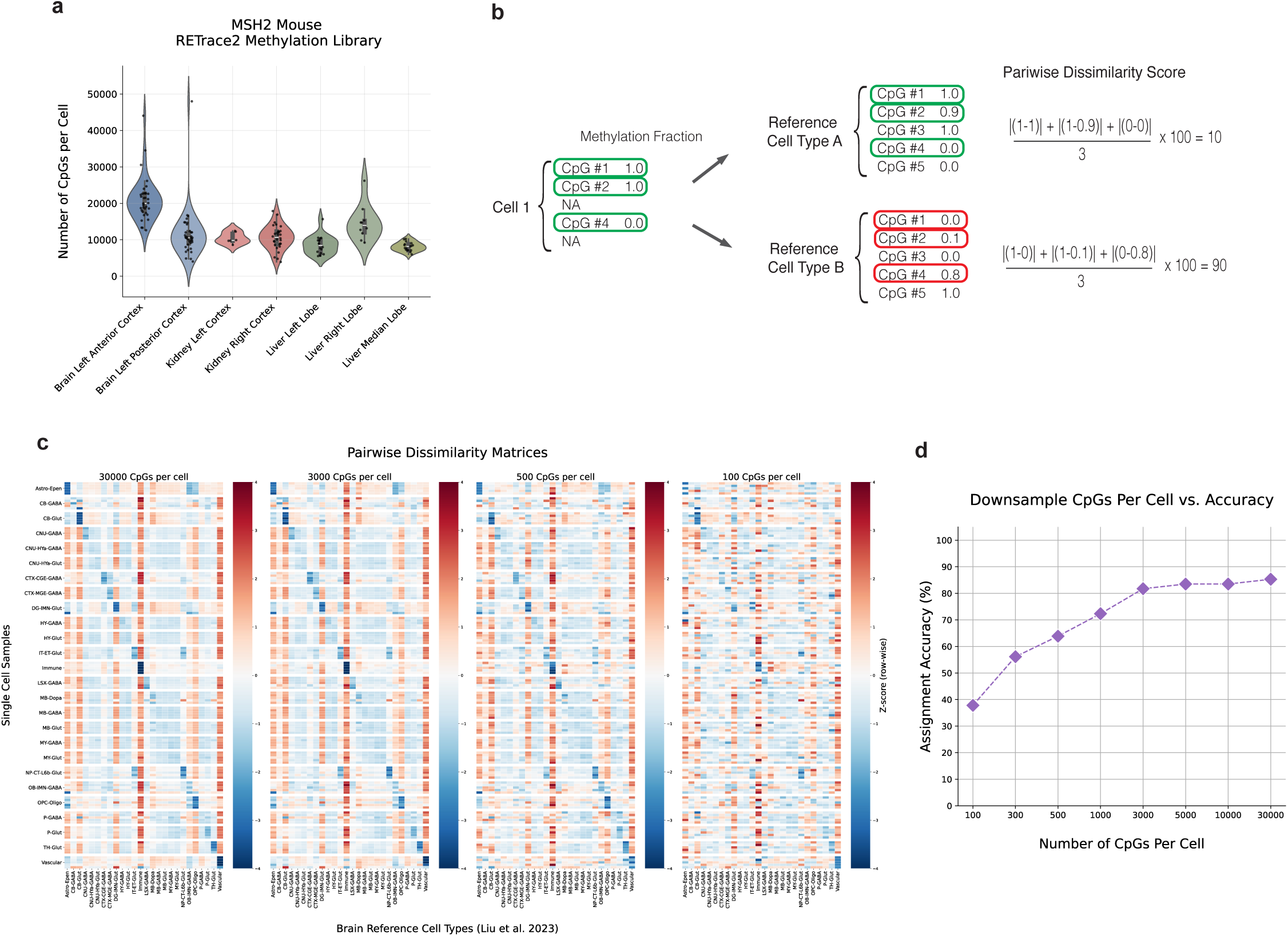
Simultaneous inference of cell type from sparse single-cell methylation profiles. **a,** Number of CpGs per cell for each tissue in the MSH2 KO/KO mouse methylation library. Each dot represents a single cell. **b,** Schematic illustrating the calculation of the pairwise dissimilarity score. **c,** Pairwise dissimilarity matrix heatmaps (z-score transformed row-wise) demonstrating the effect of computationally downsampling the number of CpGs per cell from 30,000 to 100. **d,** Cell type assignment accuracy as a function of the number of CpGs per cell, with the z-score threshold set to −1.1.

To infer cell types, we mapped our single-cell data to a reference methylome atlas by calculating a pairwise dissimilarity score, representing the mean absolute difference in methylation fractions across shared CpG sites (Fig. 5b), where a lower score indicates higher similarity. We assigned cell identities by normalizing these scores (z-score) to determine the best match, classifying cells as “unassigned” if they failed to meet a significance threshold or lacked a corresponding reference (see Methods).

We validated the robustness of this approach by simulating sparse profiles using data from the adult mouse brain single-cell methylome atlas^45^. To accurately mimic the specific coverage profile of the RETrace2 assay, we first intersected the reference data with the CpG sites captured in our experiment (reflecting the MspI restriction bias) and then computationally downsampled the CpG count per cell. Visually, serial downsampling of pairwise dissimilarity heatmaps reveals a persistent diagonal pattern, confirming that the biological signal remains distinguishable even with reduced CpG numbers (Fig. 5c). Quantitatively, we found that assignment accuracy remained high (∼81%) even with as few as 3,000 CpGs per cell (Fig. 5d), which we subsequently set as our minimum quality control filter. Applying this pipeline to our *in vivo* dataset, we successfully assigned cell types to the single cells on the phylogenetic tree (Fig. 4b, inner ring), demonstrating the method’s ability to simultaneously resolve a cell’s developmental history and its terminal identity.

## Discussion

Here, we present RETrace2, a dual-omic technology that significantly advances retrospective single-cell lineage tracing by addressing the limitations of previous methods—low marker information content and high technical noise. Our first contribution is the identification of homopolymers as superior lineage markers. We demonstrate that these mononucleotide repeats are approximately 1.9-fold more informative for lineage reconstruction than the di- to hexa-nucleotide repeats used in earlier iterations. We paired this discovery with comprehensive technical optimizations, including a vastly expanded probe set and a systematic reduction in artifacts.

Collectively, these enhancements resulted in a ∼21-fold increase in the median number of markers captured per cell and, importantly, a ∼98-fold increase in the number of shared markers per cell pair available for lineage construction. Together, these improvements allow us to harness the power of these highly mutable repeats, transforming them from a theoretically attractive marker into a practical tool.

We rigorously benchmarked this new approach using *in vitro* cell culture models with known ground-truth lineages. This validation confirmed RETrace2’s ability to resolve lineage at ∼60 cell divisions in microsatellite-stable cells, overcoming the limitation of its predecessor and establishing its potential for tracing normal, non-cancerous tissues. In microsatellite-unstable systems, such as the HCT116 cell line and the *Msh2*-deficient mouse model used here, this performance projects to a resolution of fewer than five cell divisions, offering the granularity required to reconstruct detailed developmental histories.

We next assessed the capability of RETrace2 in a complex *in vivo* setting by reconstructing a continuous, multi-organ lineage tree spanning 152 cells from the brain, kidney, and liver of an *Msh2*-deficient mouse. This high-resolution phylogeny revealed biologically meaningful clonal structures, confirming that organ development is polyclonal and characterized by strong tissue- and organ-level clustering alongside the coexistence of tissue-biased and multipotent early progenitor clades, aligning with recent findings from prospective LoxCode barcoding^8^ and retrospective human phylogenies^24^. In addition, we simultaneously inferred cell identity from sparse methylation data obtained from the same single cells, demonstrating the ability to capture both lineage and cell type information.

Our data also offer a glimpse into the timing of initial lineage restriction. We first observed significant clonal asymmetry at the root of the tree (Fig. 4b), where a dominant clade separates from a minor one (approximate 90:10 split), highlighting the stochastic nature of early progenitor contribution. This high degree of asymmetry at the earliest stages of development, indicative of an early cellular bottleneck, is a recurring feature observed in human retrospective studies using somatic SNVs^21,23,24^. Beyond this initial divergence, the major topological splits occur within the first 5–6 branching generations (early blastocyst), with lineages spanning all three germ layers. Most notably, the observation that one-third of these early-diverging clades exhibit tissue bias—while the others remain multipotent—challenges the view of rigid, binary lineage steps. Instead, our findings support a “competition model” of development^46^. In this framework, multipotency is not a passive state, but a period of active multilineage priming where competing gene programs coexist. The “multipotent” clades in our tree likely capture the history of these primed progenitors, while the “biased” clades reflect early symmetry-breaking events that precede gastrulation. These observations align with recent phylogenetic studies showing that lineage bias often emerges earlier than the traditional morphological markers of gastrulation would suggest^20,21,24,47^.

While the recent pace of innovation in retrospective lineage tracing is encouraging, the field still faces challenges in achieving the resolution required to map complete developmental pathways. Current methods each have distinct trade-offs: mitochondrial DNA variants are accessible via standard single-cell transcriptomic or chromatin assays but are subject to complex inheritance dynamics driven by heteroplasmy^33^ and the potential for horizontal transfer between cells^48^; DNA methylation provides a dense, single-modality source for both lineage and identity but can be confounded by cell-type specific regulatory programs or transient cell states induced by environmental factors (e.g., inflammation, stress, oxygen levels)^35^. This study suggests that microsatellites represent a promising marker type for lineage tracing, being stable, permanent markers with high somatic mutation rates. The >1 million microsatellites present in the human genome provide a theoretical yield of ∼10 new mutations per cell division^37,38^, which in principle is sufficient to resolve all single cell divisions. RETrace2, by integrating the informative homopolymer markers and optimized protocols to acquire ∼98-fold more shared markers with lower errors, represents a significant step toward harnessing this vast theoretical potential and ultimately reconstructing mammalian lineage trees of development, disease and aging in detail.

Future technical developments include scaling the platform and further increasing its resolution and accuracy. By implementing nanodispensing automation, 384-well plate formats, and increased multiplexing capacity, we anticipate reducing per-cell costs from approximately $36 to under $10, making large-scale organism-wide studies economically feasible. We also envision improving data quality by adopting alternative library preparation methods, such as isothermal amplification^49,50^, which can minimize allele dropout and amplification artifacts. Finally, a significant limitation of our current framework arises from the use of unphased short reads, which prevents the definitive distinction between maternal and paternal alleles required for precise genetic distance calculations. Addressing this challenge—potentially through long-read sequencing or advanced phasing algorithms—represents a logical next step for further improving the resolution of RETrace2.

## Methods

### Cell culture and ground-truth lineage tree construction

The HCT116 colorectal carcinoma cell line (Coriell Institute) was cultured in McCoy’s 5A media (ATCC) supplemented with 15% FBS and 1% penicillin-streptomycin (Thermo Fisher Scientific). The 3T3-L1 embryonic fibroblast cell line (ATCC; CL-173) was cultured in DMEM (Thermo Fisher Scientific) supplemented with 10% Bovine Growth Serum (iXCells Biotechnologies) and 1% penicillin-streptomycin. Ground-truth lineage trees were constructed by adopting an established validation strategy based on single-cell expansion^28,29^. Briefly, single cells were FACS-sorted into 96-well plates containing 25% conditioned medium. Clones were expanded by serial passaging into 24-well, 6-well, and T-75 flasks upon reaching 80-90% confluency, a process taking approximately 3-4 weeks. Cells were detached using TrypLE (Thermo Fisher Scientific), neutralized with complete medium, and centrifuged at 200g for 3 minutes. Each expansion generation represented approximately 21-23 cell divisions, yielding ∼2-8 million cells. To create branching points, fully expanded clones were re-subjected to the single-cell sorting and expansion protocol. Aliquots from each generation were cryopreserved in complete medium with 10% DMSO.

### Synthetic oligonucleotide model

All synthetic oligonucleotides for artifact benchmarking were ordered as 4nmol Ultramer DNA oligos (Integrated DNA Technologies; Supplementary Table 2). We used two distinct libraries, both containing a 20 bp UMI and a 10 bp library index for demultiplexing. The first library, designed for polymerase fidelity comparison, consisted of a pool of five oligonucleotides with different microsatellite sequences: three mononucleotide repeats (15xA, 20xA, 30xA) and two dinucleotide repeats (10xAC, 15xAC). The library index identified the ground-truth microsatellite sequence. This library was amplified separately using four high-fidelity DNA polymerases following manufacturers’ protocols: KAPA HiFi HotStart (Roche, #KK2501), Q5 High-Fidelity DNA Polymerase (New England Biolabs, #M0491), SeqAmp DNA Polymerase (Takara Bio, #638509), and PrimeSTAR Max Polymerase (Takara Bio, #R045A). Products were pooled and sequenced on an Illumina MiSeq platform (150 bp paired-end). Raw FASTQ files were processed using a custom Python script (fastq_microsatellite_extractor.py) to extract UMI, library index, and microsatellite sequences. After filtering for perfect library indices, PCR fidelity was assessed using UMI-based consensus calling. For UMI groups with ≥10X read depth, the mode (most frequent) microsatellite length was designated as the “true” sequence. Per-UMI accuracy was calculated as the percentage of reads within the group matching this mode length. Statistical comparisons were performed using a pairwise Mann-Whitney U test (KAPA HiFi as control) with Bonferroni correction (α = 0.0033). The second library, for sequencing platform accuracy comparison, contained five poly-adenine homopolymer repeats (10xA, 15xA, 20xA, 25xA, and 30xA). This library was amplified for 10 cycles using SeqAmp DNA Polymerase and split for paired-end sequencing on an Illumina NextSeq 2000 and an Element Bio AVITI. A separate custom Python script (fastq_paired_microsatellite_extractor.py) extracted UMI, library index, and homopolymer sequences (≥6 consecutive A’s) from both Read 1 (R1) and Read 2 (R2). After filtering for valid library indices, sequencing-specific accuracy was quantified as the percentage of read pairs showing perfect homopolymer length agreement between R1 and R2.

### Msh2 mouse model, tissue collection, and nuclei isolation

All animal work was approved by the Federal Ministry of Education, Science and Research, Department for Animal Testing and Genetic Engineering of Austria (license number BMWFW-66.009/0018-WF/V/3b/2017; amendment GZ-2024-0.290.102) under project leader Igor Adameyko. Heterozygous Msh2 knockout mice (Strain: B6.129P2-Msh2tm1Mak/Mmjax^42^; The Jackson Laboratory stock #42057) were paired to generate wild-type (WT/WT), heterozygous (KO/WT), and homozygous knockout (KO/KO) littermates. Tissues were collected at 6.5 weeks of age. For MSI validation, genomic DNA was extracted from small brain, kidney, and liver pieces via Proteinase K digestion. MSI status was determined by PCR amplification of four fluorescently-labeled microsatellite markers (Bat24, Bat37, Bat59, Bat64; Supplementary Table 2) followed by capillary electrophoresis (Eton Bioscience Inc). For RETrace2 library preparation, specific tissue regions were dissected: a ∼10 mg piece (∼3×3×2 mm) from a liver lobe tip; ∼40 mg of outer kidney cortex; and for the brain, one hemisphere was sagittally bisected, its cortex sectioned into four equal segments along the anterior-posterior axis, and the most anterior (1/4) and posterior (1/4) segments were used. Nuclei were isolated using the Nuclei Isolation Kit (10x Genomics), stained with Hoechst 33342 (Thermo Fisher Scientific), and filtered prior to FACS sorting.

### Single-cell and nuclei sorting and lysis

Single cells or nuclei were sorted into 96-well plates containing 5.34 µL of lysis buffer per well using a BD Aria Fusion Sorter. The lysis buffer consisted of 20 mM Tris and 2 mM EDTA (Sigma-Aldrich), 20 mM KCl (Sigma-Aldrich), 0.3% Triton X-100 (Sigma-Aldrich), 1 mg/ml protease (Qiagen), and 60 fg unmethylated Lambda DNA (Promega). Plates were immediately centrifuged at 4820g for 5 minutes at 4°C. To ensure complete lysis, plates were incubated at 50°C for 3 hours, followed by 30 minutes at 75°C to inactivate the protease, then centrifuged and stored at −80°C.

### Custom probe design and production

Homopolymer target regions (≥10 bp mononucleotide repeats) were identified in mouse (mm39) and human (hg38) reference genomes using a custom C script (find_mono-nucleotide.c). MseI digestion sites were computationally predicted using RE_selector-v1.1.py to simulate genome-wide fragmentation and select fragments compatible with 250 bp reads. Fragments were filtered to be ≥120 bp in length with 10-30 bp mononucleotide repeats. Filtered sequences were aligned to reference genomes with BWA-MEM to ensure uniqueness (mapping quality ≥1). From unique loci, 120 bp capture probes were designed using the IDT xGen Custom Hyb Panel design tool (1x tiling density, ≤30 bp end gap), retaining only probes passing IDT quality filters. Universal adapters were added to 5’ and 3’ ends, resulting in 180 bp oligonucleotides synthesized by Twist Bioscience.

Custom probe panels were generated from these oligo pools. ssDNA oligos were converted to dsDNA by PCR using primers T7_prodPCR_1 and T7_prodPCR_2 (Supplementary Table 2) with KAPA HiFi Hotstart ReadyMix (Roche) and purified with a DNA Clean & Concentrator-5 kit (Zymo Research). Purified dsDNA served as a template for in vitro transcription using T7 RNA Polymerase (New England Biolabs) for 18 hours at 37°C. RNA probes were purified using an ssDNA/RNA Clean & Concentrator kit (Zymo Research), then reverse transcribed to biotinylated ssDNA using Maxima H Minus Reverse Transcriptase (Thermo Fisher Scientific) and a biotinylated primer (T7_RT_primer; Supplementary Table 2). The RNA template was removed by RNase H digestion (New England Biolabs), and final biotinylated ssDNA probes were purified using the ssDNA/RNA Clean & Concentrator kit. Concentration and quality were assessed by Qubit fluorometer and TapeStation.

### RETrace2 library preparation

Following lysis, single-cell gDNA was digested with MseI (Thermo Fisher Scientific) and MspI (New England Biolabs) and simultaneously A-tailed using Klenow fragment (exo-) (Thermo Fisher Scientific). NEBNext EM-seq adapters (New England Biolabs) were ligated using the NEBNext Ultra II Ligation Module (New England Biolabs). The library was amplified for 20 cycles with a single primer (Bst_RCA_Primer_v4; Supplementary Table 2) using KAPA HiFi DNA Polymerase (Roche). The product was split: 20 µL for methylation and 5 µL for microsatellite library preparation.

The methylation aliquot underwent bisulfite conversion using the EZ-96 DNA Methylation-Direct MagPrep kit (Zymo Research), following a modification of the scRRBS protocol^44^. Converted DNA was amplified for 32 cycles with KAPA HiFi Uracil+ ReadyMix (Roche) and dual-indexed primers for MspI-digested fragments (Supplementary Table 2). The final methylation library was purified using 0.8x AMPure XP beads (Beckman Coulter) with two 80% ethanol washes.

The microsatellite aliquot was amplified for 6 cycles with SeqAmp DNA Polymerase (Takara Bio) and dual-indexed primers for MseI-digested fragments (Supplementary Table 2), then purified with 0.8x AMPure XP beads. For enrichment, 500-2000 ng of library was hybridized for 16-18 hours to a custom biotinylated homopolymer probe panel. Probe amount was scaled with complexity: ∼3 pmol for small sets (<24k probes) and ∼26 pmol for the 169k mouse panel. Hybridization used the xGen Hybridization and Wash Kit (IDT) with xGen Universal Blockers (IDT). Targets were captured on Dynabeads M-270 Streptavidin, washed, and amplified on-bead for 5-7 cycles using SeqAmp DNA Polymerase supplemented with 1.5 mM MgCl2 (final 2.5 mM). The final library was purified with 1.2x AMPure XP beads. Microsatellite and methylation libraries were pooled at a 9:1 ratio (targeting ∼9M and ∼1M reads per cell, respectively) and sequenced on an Element Bio AVITI with 10% PhiX spike-in (Read 1: 250 bp; Index 1 & 2: 8 bp).

### Microsatellite data processing

Single-cell sequencing data were processed using the RETrace2 pipeline, a Nextflow-based bioinformatics workflow for dual-omic lineage tracing and cell type inference. For microsatellite libraries, raw FASTQ files underwent quality control with FastQC (v0.12.1), followed by adapter trimming with Trim-galore (v0.6.10) using quality threshold 30 and minimum read length 36 bp. Trimmed reads were aligned to the reference genome using BWA-MEM (v0.7.19) with duplicate reads retained, followed by coordinate sorting and indexing with SAMtools (v1.21). Samples were filtered based on the number of captured microsatellite loci: a minimum of 500 targets per cell was required for the HCT116 cell line dataset processed with the smaller test probe set, while 10,000 targets per cell was required for the full probe set analyses. Microsatellite genotypes were called using HipSTR with quality filtering parameters: minimum 10 reads per locus and maximum stutter ratio 1.0. For the full probe set data, a minimum quality score of 0.9 was applied to ensure high-confidence genotype calls. For the HCT116 test dataset, this filter was disabled (set to 0) to prevent the pre-filtering of any targets, a necessary step for the subsequent downsampling analysis to ensure an unbiased comparison of the probe set’s full informative potential. The VCF output was parsed to extract allele information, grouping alleles with consecutive repeat lengths into allele classes for phylogenetic analysis. Pairwise genetic distances between cells were calculated using a minimum comparison strategy with binary distance metrics (minComp_EqorNot), which treats alleles as either matching (distance = 0) or different (distance = 1) while accounting for potential allelic dropout by comparing only the minimum number of alleles present between samples. Phylogenetic trees were reconstructed using the neighbor-joining algorithm^51^ implemented in scikit-bio (v0.6.3) with midpoint rooting.

### Methylation data processing and cell type inference

Methylation libraries were processed with FastQC for quality control, followed by RRBS-specific adapter trimming using Trim-galore (quality threshold 20, minimum read length 36 bp, RRBS mode enabled). Trimmed reads were aligned and methylation calls generated using the methylpy (v1.4.7) single-end pipeline with minimum base quality score 30. A minimum of 3,000 CpGs per cell was required for analysis. Brain cell type reference methylomes were constructed from single-cell ALLC files from Liu et al. (2023)^45^. For each major cell type, up to 3,000 individual cells were aggregated into pseudobulk references by summing methylated and total read counts at each CpG site. To infer cell types, pairwise methylation dissimilarity scores were calculated between single-cell and reference methylomes using a custom Python script. For each cytosine site, the methylation fraction was calculated as the number of methylated reads divided by the total reads. For each single cell-reference pair, the dissimilarity score was calculated as the mean absolute difference in methylation fractions across all shared CpG sites (minimum 1 read per site required). Only comparisons with at least 100 shared CpG sites were considered valid. For cell type assignment, we first computed row-wise z-scores for each single cell across all reference dissimilarity values using the formula: z-score = (dissimilarity - row mean) / row standard deviation. This normalization accounts for cell-to-cell variation in overall methylation patterns. Each cell was then assigned to the reference cell type with the minimum z-score, provided the z-score was below a threshold of −1.1, indicating significantly lower dissimilarity compared to other reference types. Cells failing to meet this threshold for any reference cell type, or those from tissues for which no reference methylome was available, were classified as ‘unassigned’.

### Tree accuracy calculation

For the cell culture model with ground-truth, tree accuracy was quantified by comparing the topology of the reconstructed tree to the known ground-truth lineage. This was done by evaluating all possible three-cell combinations, or “triplets,” where at least two of the three cells originated from different clones. For each triplet, the pair of cells separated by the greatest distance from their most recent common ancestor (MRCA) was identified in both the reconstructed tree and the ground-truth tree. A triplet was scored as “correct” if this most-distant pair was identical in both trees. The overall accuracy represents the percentage of all evaluated triplets that were scored as correct. A randomly generated tree is expected to have a baseline accuracy of ∼33%.

### Downsampling analysis and linear regression

To evaluate the information content of different microsatellite types, we performed a computational downsampling analysis. For a given number of targets (e.g., 100, 150, 200), we first identified all cells in the experiment that had captured at least that many targets. From this subset of cells, we then created 30 iterations by randomly sampling the specified number of targets from each cell’s total captured targets. Only iterations containing at least 30 cells were included for robust evaluation. Linear regression was then performed on the complete set of data points (tree accuracy vs. number of targets across all iterations) to calculate the slope, R^2^ value, and P-value. To test for a significant difference between the slopes of the regression lines, an Analysis of Covariance (ANCOVA) was performed using the statsmodels library in Python.

### Permutation tests for tissue clustering

The statistical significance of tissue clustering in the phylogenetic tree was evaluated using two permutation tests. The test statistic for both was the percentage of sibling tips (cells sharing the same immediate parent node with a cophenetic distance ≤ 2) that originated from the same tissue. The first test assessed the null hypothesis that the observed tissue clustering is no better than random label assignment. This was done by calculating the test statistic on the fixed experimental tree after randomly shuffling the tissue labels among all cells (1,000 permutations). The second, complementary test assesses whether the microsatellite genotypes themselves contain the biological signal that drives the tree’s structure. For this test, the genotype information was shuffled across samples independently for each locus, a new phylogenetic tree was reconstructed from the permuted genotypes, and the test statistic was calculated. This process was repeated 100 times. For both tests, an empirical p-value was calculated as the proportion of permutations that resulted in a test statistic greater than or equal to the experimentally observed value.

### Phylogenetic clade and tissue bias analysis

To quantitatively assess tissue-specific enrichment, clades (subtrees) were first identified from the reconstructed phylogenetic tree using a recursive traversal algorithm. Major clades were defined as non-overlapping subtrees containing between 10 and 20 cells. For each identified clade, tissue bias was evaluated using a one-sided hypergeometric test to determine if any of the seven granular tissue subtypes were significantly over-represented compared to their overall frequency in the full dataset. P-values were corrected for multiple comparisons across the seven tissues using the Benjamini-Hochberg False Discovery Rate (FDR) procedure. A clade was classified as having a “strong bias” if a tissue was enriched with an FDR < 0.05 and a fold-change enrichment of at least 2.0. Clades with no tissue meeting an FDR < 0.2 were classified as “multipotent”.

### K-Nearest Neighbor (KNN) analysis

To characterize the clonal neighborhood composition, we performed a KNN analysis. First, pairwise genetic distances between all cells were calculated. These raw distances were then adjusted for total divergence to account for different mutation rates across lineages by computing the Q-matrix, which uses the same logic as the neighbor-joining algorithm, to normalize distances based on each cell’s total divergence. For each cell, we identified its 15 nearest neighbors based on this divergence-adjusted distance matrix. The tissue types of these neighbors were counted to generate a neighborhood composition profile for each cell. To correct for differences in tissue abundance, these counts were normalized to yield relative enrichment values. Tissues were then compared using Pearson correlation based on their enrichment profiles, and hierarchically clustered.

## Data availability

All raw sequencing data (microsatellite and methylation libraries) generated in this study have been deposited in the NCBI Sequence Read Archive (SRA) under BioProject accession number PRJNA1377203. All processed data files, including microsatellite genotypes, methylation calls, phylogenetic trees, and custom code used for analysis, are publicly available on Zenodo at https://doi.org/10.5281/zenodo.17945194.

## Code availability

RETrace2 nextflow pipeline is available on GitHub (https://github.com/tonypincheng/RETrace2). Jupyter notebooks to reproduce the analyses and figures presented in this study are also available in the repository’s *notebooks* directory.

## Protocol availability

A detailed, step-by-step protocol for the RETrace2 library preparation is publicly available at protocols.io: dx.doi.org/10.17504/protocols.io.yxmvmb12bg3p/v1.

## Supplementary Information

**Extended Data Fig. 1.**
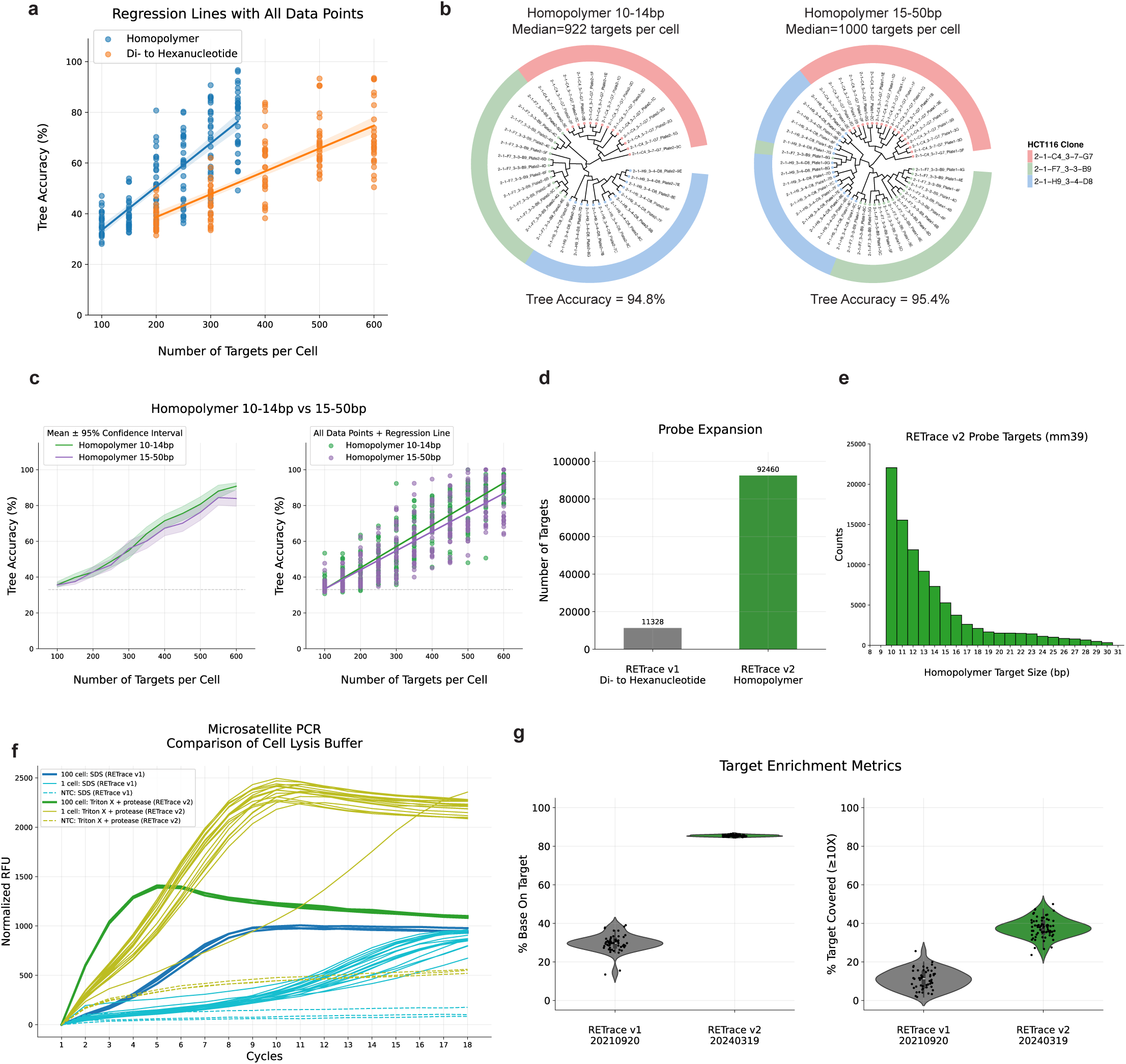
Comparison of microsatellite informativeness and protocol optimizations. **a,** Linear regression analysis of phylogenetic tree accuracy versus the number of targets per cell for homopolymer (n=180 data points) and di- to hexanucleotide (n=150 data points) datasets. **b,** Reconstructed phylogenetic trees for HCT116 cells using probes targeting short (10-14 bp; n=41 cells) versus long (15-50 bp; n=52 cells) homopolymers. **c,** Quantitative comparison of tree accuracy for short versus long homopolymers, shown as mean ± 95% confidence interval across 30 random downsampling iterations (left) and as a linear regression of all data points (right; n=330 per group; Slopes: 0.1186 vs 0.1068; R²: 0.7966 vs 0.7234; ANCOVA p=0.016). **d,** Number of targets in the RETrace v1 versus the expanded v2 probe sets. **e,** Size distribution of homopolymer targets within the RETrace2 mouse probe set. **f,** qPCR amplification curves from 3T3L1 cells, comparing the cell lysis efficiency of the RETrace v1 (SDS-based) and v2 (Trypsin+protease-based) protocols. **g,** Target enrichment metrics for pre-filter RETrace v1 (20210920 di to hexanucleotide library; n=54) and v2 (20240319 library; n=72). Each dot represents a single cell.

**Extended Data Fig. 2.**
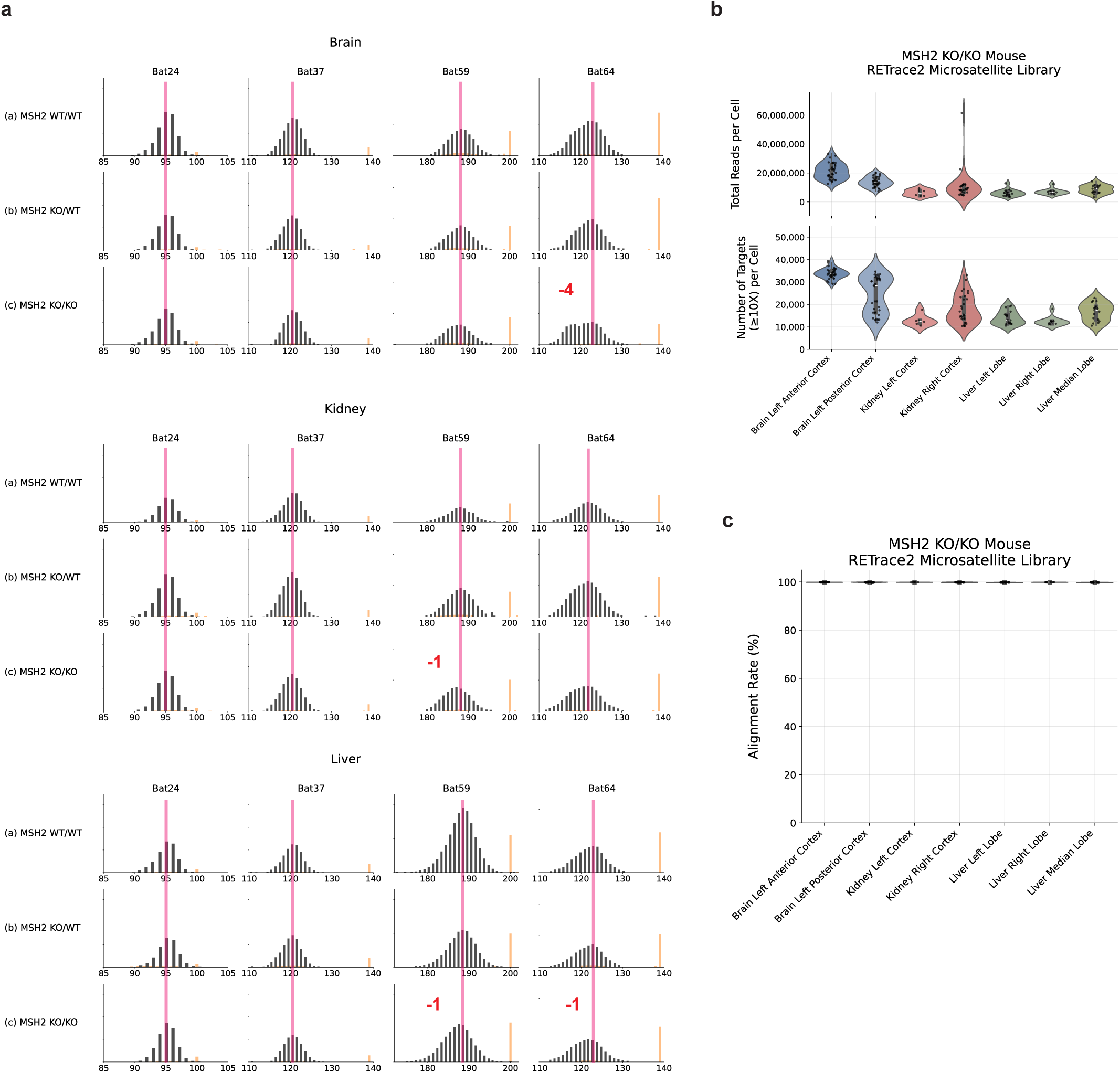
Validation of microsatellite instability in the Msh2 mouse model and microsatellite library QC. **a,** Capillary electrophoresis fragment analysis confirming microsatellite instability (MSI) across three tissues (brain, top; kidney, middle; liver, bottom) from littermates with wild-type (WT/WT), heterozygous (KO/WT), and homozygous knockout (KO/KO) Msh2 genotypes. Four distinct homopolymer markers (columns) were tested. The KO/KO mouse shows altered fragment sizes, with red numbers indicating the size shift in base pairs (bp) relative to the WT/WT control (red bars), indicative of MSI. Yellow bars represent internal size standards used for fragment analysis. **b,** Quality control metrics for the RETrace2 microsatellite library, showing the distribution of total reads per cell (top) and the number of targets covered at ≥10X depth per cell (bottom) for each tissue. **c,** BWA alignment rate for the RETrace2 microsatellite library across all seven dissected tissues.

**Extended Data Fig. 3.**
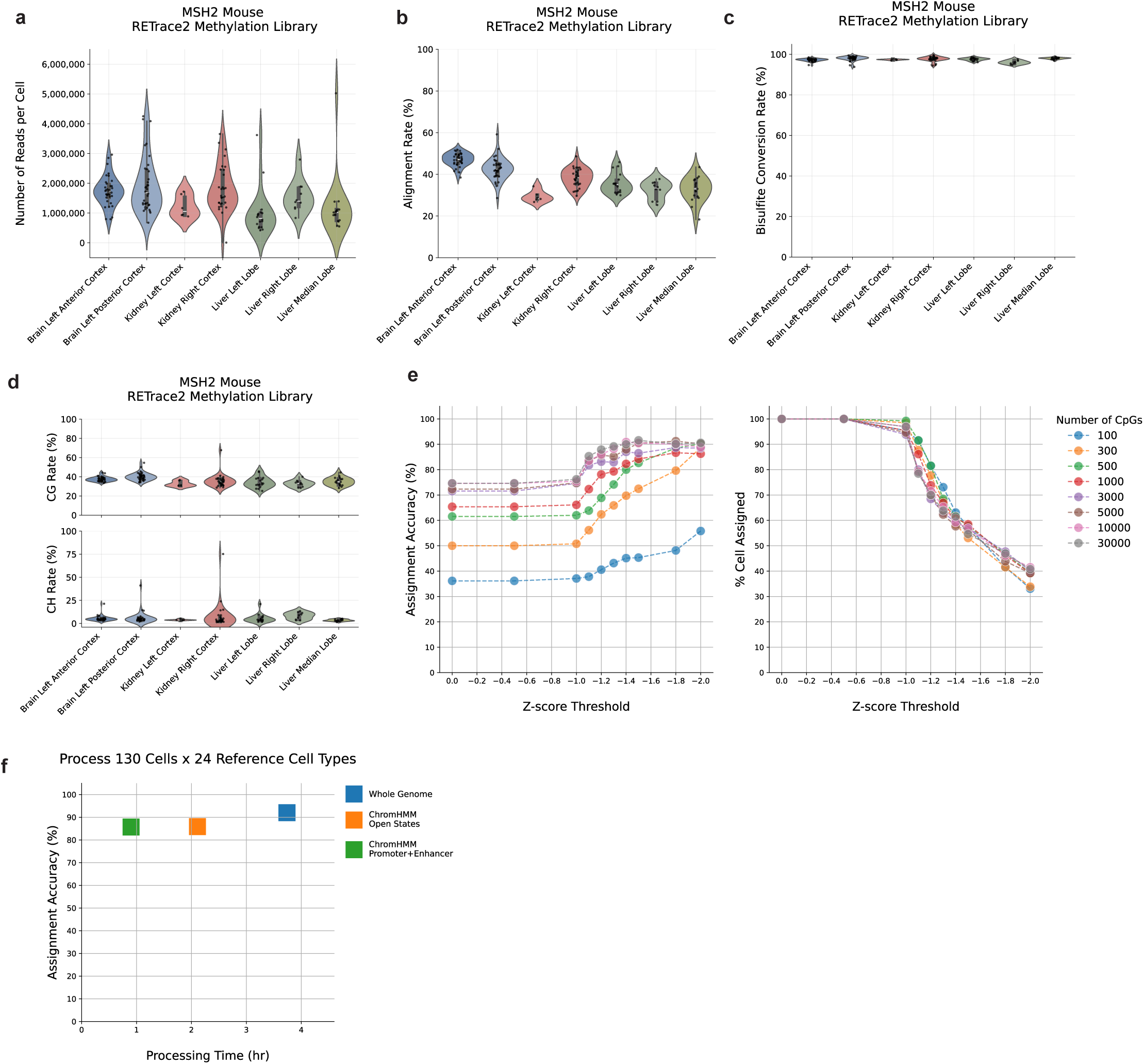
RETrace2 methylation library QC and computational pipeline validation. **a-d,** Quality control metrics for the MSH2 KO/KO mouse methylation library, showing total reads (a), alignment rate (b), bisulfite conversion rate (c), CG and CH methylation rates (d). Each dot represents a single cell. **e,** Plots calibrating the z-score threshold for cell assignment. The trade-off between assignment accuracy (left) and the percentage of cells assigned (right) is shown across different z-score thresholds, guiding the selection of an optimal parameter. **f,** Analysis of computational efficiency. Comparison of cell type assignment accuracy and processing time when mapping to the whole genome versus more restricted, biologically relevant genomic regions (ChromHMM-defined open chromatin states or promoter and enhancer regions). The analysis demonstrates that mapping to these focused regions substantially reduces computational time with minimal impact on accuracy.

## Acknowledgements

We thank Christopher Jen-Yue Wei for laying the foundation of RETrace v1 and for proposing the potential of homopolymers as lineage markers in that work. We thank the Altos Labs Genomic Core for library sequencing. We thank the team at Element Bio for their valuable collaboration and technical support in testing the AVITI sequencing platform. We are also grateful to Kimberly Conklin for lab management and logistical support, and to all members of the Zhang lab for their helpful discussions and feedback on the manuscript.

## Author Contributions

P.C.C. conceptualized and developed the RETrace2 method, performed the *in vitro* and *in vivo* experiments, analyzed all lineage tracing and methylation data, and wrote the manuscript. D.K., P. Kameneva, and I.A. generated the Msh2 mouse model, advised on tissue selection, and collected tissues for the experiments. C.F. performed the single cell and nuclei sorting. P.V.K. provided conceptual advice on statistical analyses, including the permutation test. K.Z. conceptualized the study, provided guidance and resources, and supervised all aspects of the work. All authors read and approved the final manuscript.

