## Supplementary Fig. 1 for "High-resolution retrospective single cell lineage tracing with mutable homopolymers"

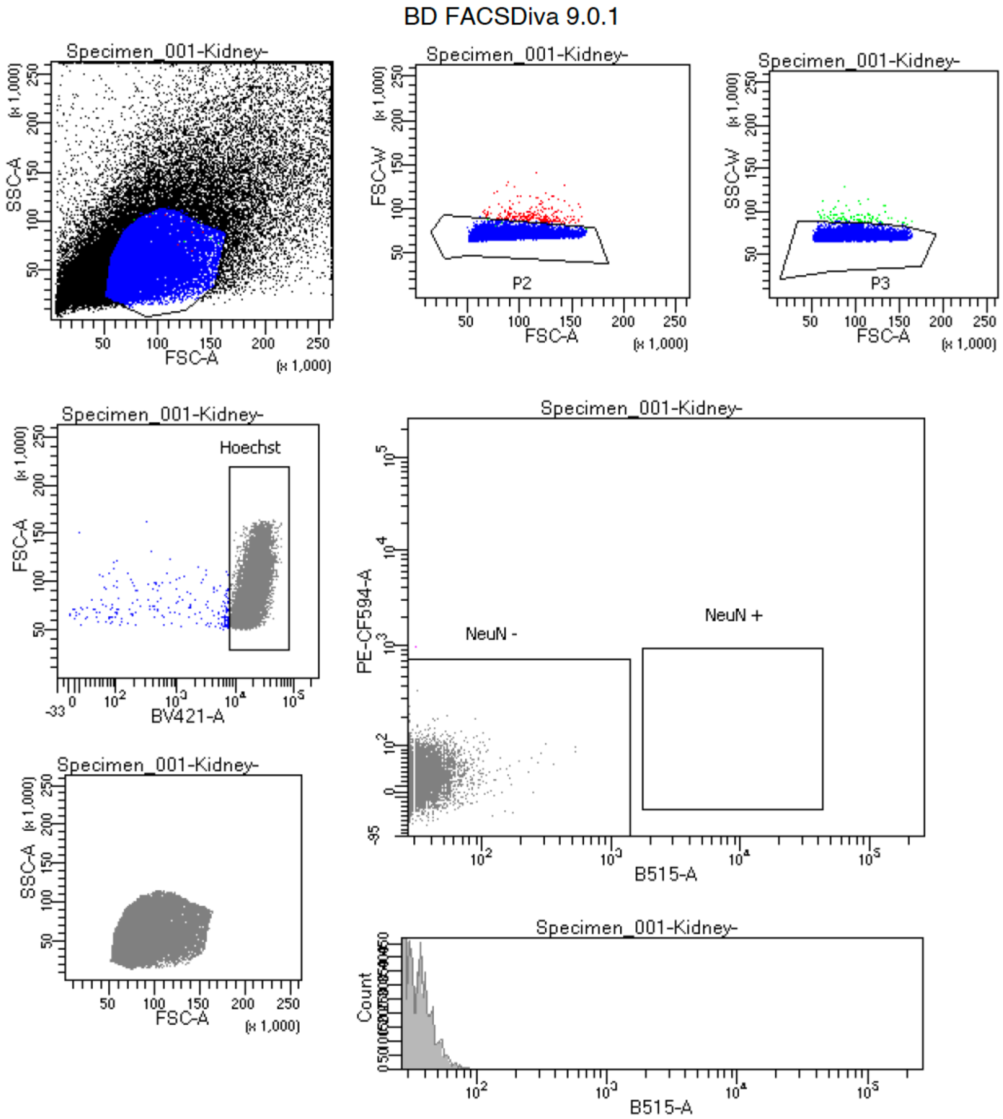


Tube: Kidney-

| Population | #Events | %Parent | %Total |
|---|---|---|---|
| All Events | 41,964 | #### | 100.0 |
| P1 | 15,880 | 37.8 | 37.8 |
| P2 | 15,653 | 98.6 | 37.3 |
| P3 | 15,558 | 99.4 | 37.1 |
| Hoechst | 15,313 | 98.4 | 36.5 |
| NeuN - | 15,312 | 100.0 | 36.5 |
| NeuN + | 0 | 0.0 | 0.0 |

**Supplementary Fig. 1 | Representative FACS gating strategy for single-nuclei isolation.** Gating hierarchy for sorting single nuclei from MSH2 knockout mouse tissues (representative Kidney sample shown). Nuclei were first identified by light scatter properties (FSC-A vs. SSC-A) to exclude debris (P1). Doublets and aggregates were removed using pulse-width geometry gating on FSC-W vs. FSC-A (P2) and SSC-W vs. SSC-A (P3). Singlet nuclei were verified by Hoechst 33342 fluorescence (BV421-A) prior to sorting into lysis buffer.
